# Preclinical immunoPET imaging of glioblastoma-infiltrating myeloid cells using Zirconium-89-labeled anti-CD11b antibody

**DOI:** 10.1101/614511

**Authors:** Shubhanchi Nigam, Lauren McCarl, Rajeev Kumar, Robert S. Edinger, Brenda F. Kurland, Carolyn J. Anderson, Ashok Panigrahy, Gary Kohanbash, W. Barry Edwards

**Author notes:** Authors contributed equally (Shubhanchi Nigam, Lauren McCarl). co-corresponding author Wilson B. Edwards, PhD, Gary Kohanbash, PhD.

## Abstract

**Purpose:** Glioblastoma is a lethal brain tumor, heavily infiltrated by tumor-associated myeloid cells (TAMCs). TAMCs are emerging as a promising therapeutic target as they suppress antitumor immune responses and promote tumor cell growth. Quantifying TAMCs using non-invasive immunoPET could facilitate patient stratification for TAMC-targeted treatments and monitoring of treatment efficacy. As TAMCs uniformly express the cell surface marker, integrin CD11b, we evaluated a ^89^Zr-labeled anti-CD11b antibody for non-invasive imaging of TAMCs in a syngeneic orthotopic mouse glioma model.

**Procedures:** A human/mouse cross-reactive anti-CD11b antibody (clone M1/70) was conjugated to a DFO chelator and radiolabeled with Zr-89. PET/CT and biodistribution with or without a blocking dose of anti-CD11b Ab were performed 72 hours post-injection of ^89^Zr-anti-CD11b Ab in mice bearing established orthotopic syngeneic GL261 gliomas. Flow cytometry and immunohistochemistry of dissected GL261 tumors were conducted to confirm the presence of CD11b^+^ TAMCs.

**Results:** Significant uptake of ^89^Zr-anti-CD11b Ab was detected at the tumor site (SUVmean = 2.60 ± 0.24) compared with the contralateral hemisphere (SUVmean = 0.6 ± 0.11). Blocking with a 10-fold lower specific activity of ^89^Zr-anti-CD11b Ab markedly reduced the SUV in the right brain (SUVmean = 0.11 ± 0.06), demonstrating specificity. Spleen and lymph nodes (myeloid cell rich organs) also showed high uptake of the tracer, and biodistribution analysis correlated with the imaging results. CD11b expression within the tumor was validated using flow cytometry and immunohistochemistry, which showed high CD11b expression primarily in the tumoral hemisphere compared to the contralateral hemisphere.

**Conclusion:** These data establish that ^89^Zr-anti-CD11b Ab immunoPET targets CD11b^+^ cells (TAMCs) with high specificity in a mouse model of GBM, demonstrating the potential for non-invasive quantification of tumor infiltrating CD11b^+^ immune cells during disease progression and immunotherapy in patients with GBM.

## Introduction

Glioblastoma (GBM; also referred to as a grade IV glioma), is the most common primary malignant central nervous system tumor, with over 12,000 new cases diagnosed each year in the United States. Despite aggressive treatment with maximum safe surgical resection, radiotherapy, and chemotherapy (temozolomide), the median overall survival time for these patients is less than two years [1]. Immunotherapy may be a promising treatment option for GBM patients; however, effective immunotherapies must overcome the immune-suppressive tumor microenvironment.

Immunosuppression in GBM is largely mediated through the recruitment of T-regulatory cells [2] and tumor-associated myeloid cells (TAMCs). TAMCs primarily include two cell populations: tumor-associated macrophages (TAMs; including peripheral origin and brain-intrinsic microglia) and myeloid-derived suppressor cells (MDSCs) [3]. TAMCs account for up to 40% of a glioma’s cellular mass and play an important role in tumor progression, immunosuppression, and resistance to immunotherapy [4-6]. Numerous preclinical studies by us and others have demonstrated that targeting TAMs and MDSCs can prolong survival of mice with gliomas [5, 7-8] and that these cells are associated with poor prognosis in high-grade gliomas [9]. Since TAMCs impede natural and immunotherapy-driven anti-tumor responses, they are a high-priority and promising therapeutic target currently being evaluated in clinical trials [7-8, 10]. Accordingly, quantifying these immunosuppressive cells will enable stratification of patients and monitoring of treatment efficacy [10-14].

Current methods to monitor tumor-infiltrating immune cells include multiplexed flow cytometry, proteomics, genomics technologies (*e.g.* single cell sequencing and mass cytometry), and histology on surgically removed tumors [15-18]. While these approaches directly measure TAMC levels, they lack the temporal and spatial resolution of non-invasive diagnostic imaging. Although MRI is a powerful diagnostic tool for GBM, it lacks the capacity to directly quantify the status of immune cell populations throughout the tumor microenvironment [19].

Positron-emission tomography (PET) provides the real-time status of tumor-infiltrating immune cells during disease progression and treatment. Using appropriate radiotracers, PET may be able to improve therapeutic responses by selecting early-phase responders and guiding the adjustment of treatment strategies. There is currently an unmet need for PET tracers to quantify the highly dynamic populations of tumor-infiltrating immune cells. To overcome this limitation, we have validated a ^89^Zr-labeled anti-human/mouse CD11b antibody for targeting both MDSCs and TAMCs in an orthotopic mouse GBM model by immunoPET. The same strategy as presented here in mouse models will ultimately be applied with anti-human CD11b antibodies in patients with GBM.

## Materials and Methods

### Reagents

All chemicals were purchased from Sigma-Aldrich Chemicals (St. Louis, MO) unless otherwise specified. Aqueous solutions were prepared using ultrapure water (resistivity, 18.2 MΩ ·cm). Para-NCS-Bz-DFO was purchased from CheMatech (Dijon, France). All antibodies for flow cytometry were purchased from BioLegend (San Diego, CA). Zirconium-89 oxalate was purchased from Washington University (St. Louis, MO).

### Instrumentation

Radiochemistry reaction progress and purity were monitored using BIOSCAN ITLC (Eckert & Ziegler, Germany) and an Agilent 1260 infinity HPLC (Agilent Technologies, Santa Clara, CA) using Superose™ 12 10/300 GL SEC column (GE Healthcare, Chicago, IL). MRI data were acquired on a Bruker ClinScan 7 T MRI (Billerica, MA). Biodistribution samples were counted using a PerkinElmer 2470 WIZARD2 Automatic Gamma Counter (Waltham, MA). PET/CT data were acquired on an Inveon Preclinical Imaging Station (Siemens Medical Solutions, Knoxville, TN). Flow cytometry studies were performed on a BD LSRII (BD Biosciences, San Jose, CA), and data were analyzed using FlowJo software (BD Biosciences). IHC and hematoxylin and eosin (H&E) images were acquired using a Leica DFC7000T microscope (Leica Microsystems Inc).

### Anti-CD11b Ab conjugation with a p-NCS-Bz-DFO chelator

A para-NCS-Bz-DFO stock solution (10 mM) was prepared in dry DMSO. A portion of the para-NCS-Bz-DFO stock (4.5 μl, 45 nmole) was added to anti-CD11b Ab (clone M1/70, 0.5 mg/500 μl, 3 nanomoles). The alkalinity was adjusted to pH∼9 by the addition of sodium carbonate (5 μl, 0.1 M Na_2_CO_3_). After 18 h (at 4°C), the reaction mixture was applied to a desalting column (MWCO 50,000) and eluted with buffer (0.5 M HEPES, pH 7.4). The fractions containing the DFO-anti-CD11b conjugate were pooled, concentrated, and stored at −20°C. Size exclusion chromatography (SEC) was performed using PBS as a mobile phase to assess the purity of the conjugate.

### Radiolabeling

Zirconium-89 oxalate (74 MBq) was added to a labeling buffer (equal volumes of 1 M HEPES (100 μL) and 1 M Na_2_CO_3_ (100 μL). Then, DFO-anti-CD11b Ab (0.2 nmol in 1M HEPES) was added to the mixture for radiolabeling (1 h, 37°C). The reaction was monitored by instant thin-layer chromatography (ITLC; BIODEX, green Ab strips, New York, NY) and purified with a PD-10 desalting column (5 mg/ml gentisic acid in saline). Final radiolabeling purity was ≥ 99% with a specific activity of 11.1GBq/μmol. Radiochemical purity (≥ 99%) was also accessed by SEC-HPLC.

### Cell line and mouse model of GBM

GL261 glioma cells were cultured in Dulbecco’s Modified Eagle Medium (DMEM) (Lonza, Walkersville, MD) supplemented with 2 mM L-glutamine (Life Technologies Corporation, Grand Island, NY), 1X Antibiotic-Antimycotic (Life Technologies Corporation), 1 mM sodium pyruvate (Lonza), 55 μM 2-mercaptoethanol (Life Technologies Corporation), 10% heat-inactivated fetal bovine serum (FBS; Thermo Fisher), 1X NEAA Mixture (Lonza), and 10 mg/mL Normocin (InvivoGen, San Diego, CA). Shortly before stereotactic injection, GL261 cells were washed twice with DPBS (Lonza), filtered through a 40 μm cell strainer, and resuspended in DPBS at a concentration of 50,000 cells/μL.

All animal studies were conducted under protocols approved by the University of Pittsburgh Institutional Animal Care and Use Committee (IACUC). For intracranial tumor cell injections, C57BL/6j mice (female, 5-8 weeks old; Jackson Labs, Bar Harbor, ME) were anesthetized with isoflurane and placed on a stereotactic frame (Kopf Instruments, Tujunga, CA). 1 × 10^5^ GL261 cells in 2 μL of DPBS were injected into the right caudate nucleus (at a coordinate from the skull position of bregma of: +1.5 mm AP, +2.5 mm ML, and −3.0 mm DV) using a micro-pump injector (World Precision Instruments, Sarasota, FL).

### *In vivo* imaging

MRI was performed on all GL261 tumor-bearing mice (15-17 days post tumor cell injection) one day before PET/CT imaging to calculate tumor volume (∼30 mm^3^). For PET/CT imaging, mice were injected intravenously with ^89^Zr-DFO-CD11b (3.7 MBq; specific activity: 11.1 GBq/μmol for experimental group; 1.11 GBq/μmol for blocking group). Mice were anesthetized with 2% isoflurane, and static PET/CT imaging (Inveon Preclinical Imaging Station (Siemens Medical Solutions, Knoxville, TN) was performed 72 h post injection. Images were co-registered and analyzed using Inveon Research Workstation (IRW) software (version 4.2, Siemens Healthcare, Germany). Region of interest analysis was guided by CT. ^89^Zr-anti-CD11b Ab uptake is presented as SUVmean.

### Flow cytometric assessment

GL261 tumor-bearing C57BL/6j mice (separate from the imaging cohorts) were euthanized and transcardially perfused with 10 ml of DPBS to remove circulating blood cells. Brains were harvested, and the GL261 tumors were excised and dissociated in Collagenase IV Cocktail (3.2 mg/mL Collagenase Type 4, 1.0 mg/mL deoxyribonuclease I, 2 mg/mL Soybean Trypsin Inhibitor [Worthington Biochemical, Lakewood, NJ]) for two 15 min intervals with gentle mixing at 37°C. The resulting tumor cell suspensions were filtered through 70 μm cell strainers. Red blood cells were subsequently lysed with ACK Lysing Buffer (Lonza), and the suspensions were filtered through 40 μm cell strainers.

For flow cytometric analysis, dissociated GL261 tumors (5 × 10^6^ cells) were pre-incubated with 1.0 μg of TruStain fcX (anti-mouse CD16/32, clone 93; BioLegend) in 100 μL of FACs Buffer (DPBS with 1% BSA) for 10 min on ice. Cells were then incubated with APC-anti-mouse CD45 (BioLegend, clone 30-F11) and PE-anti-mouse/human CD11b (BioLegend, clone M1/70) or their respective isotype controls (APC Rat IgG2b, BioLegend, clone RTK4530 and PE Rat IgG2b, BioLegend, clone RTK4530) for 30 min at 4°C. Stained cells were washed twice with FACs Buffer, fixed using Fixation Buffer (BioLegend), and resuspended in Cyto-Last Buffer (BioLegend) for cytometric analysis. Flow cytometric analysis was performed on a BD LSRII cytometer, and the data were analyzed using FlowJo software (version 10.5.3). The number of CD11b receptors on CD45^+^CD11b^+^ TAMCs was quantified using the QuantiBRITE PE Fluorescence Quantitation Kit (BD Biosciences), according to the manufacturer’s recommended protocol.

### Immunohistochemistry analysis

For immunohistochemical analysis, the PET/CT imaging and biodistribution cohort (n=6) tumors were used. Tumor ipsilateral right brain and contralateral left-brain samples were fixed in neutral phosphate-buffered 10% formalin (Fisher Scientific, Pittsburgh, PA) for 24 h. Samples were then transferred to 70% ethanol until the Zr-89 decayed (>5 half-lives). Samples were dehydrated using an EtOH gradient of 70% to 100% over 24 h, cleared with histology grade xylene (Fisher Scientific), embedded in granular paraffin, and sectioned (5 μm) on poly-lysine slides. Sections were incubated with 10% normal goat serum, and then overnight at 4°C with primary antibody. Biotinylated secondary goat anti-rabbit Ab (Abcam; dilution 1:200) was used to detect the primary Ab. The staining signal was amplified with avidin-biotin complex (Vector Laboratories, Burlingame, CA) and developed using chromogen 3-amino-9-ethylcarbazole (Sigma-Aldrich). ImageJ software (ImmunoRatio plugin) was used to quantify CD11b^+^ cells per nuclear area [20]. A histomorphologic analysis of the tumors was performed with the H&E slides.

### Statistical analysis

Right brain (tumor) and left brain (contralateral control) SUVmean were compared using a paired t-test. Experimental and blocking group SUVmean values were compared using t-tests for independent groups (unequal standard deviations). Statistical analyses were performed using GraphPad Prism 7 (GraphPad, San Diego, CA). All tests were two-sided; a p-value <0.05 was considered significant. When multiple endpoints were compared in a single analysis (experimental-blocking comparisons for organ SUVmean and biodistribution), the false discovery rate was controlled to 0.05 by the method of Benjamini and Hochberg [21].

## Results

### Radiolabeling, Stability, and Immunoreactivity

Assessment of radiochemical yields (>99%, retention time of 18 min) **(Fig. 1)** and purity of ^89^Zr-anti-CD11b showed >99% stability in PBS and mouse serum at 37°C up to 3 days. The radiolabeled antibody was mono-molecular with no signs of aggregation. In isolated splenocytes, there was 12%-15% immunoreactivity, consistent with the fraction of CD11b^+^ cells in normal mouse spleen.

**Figure 1.**
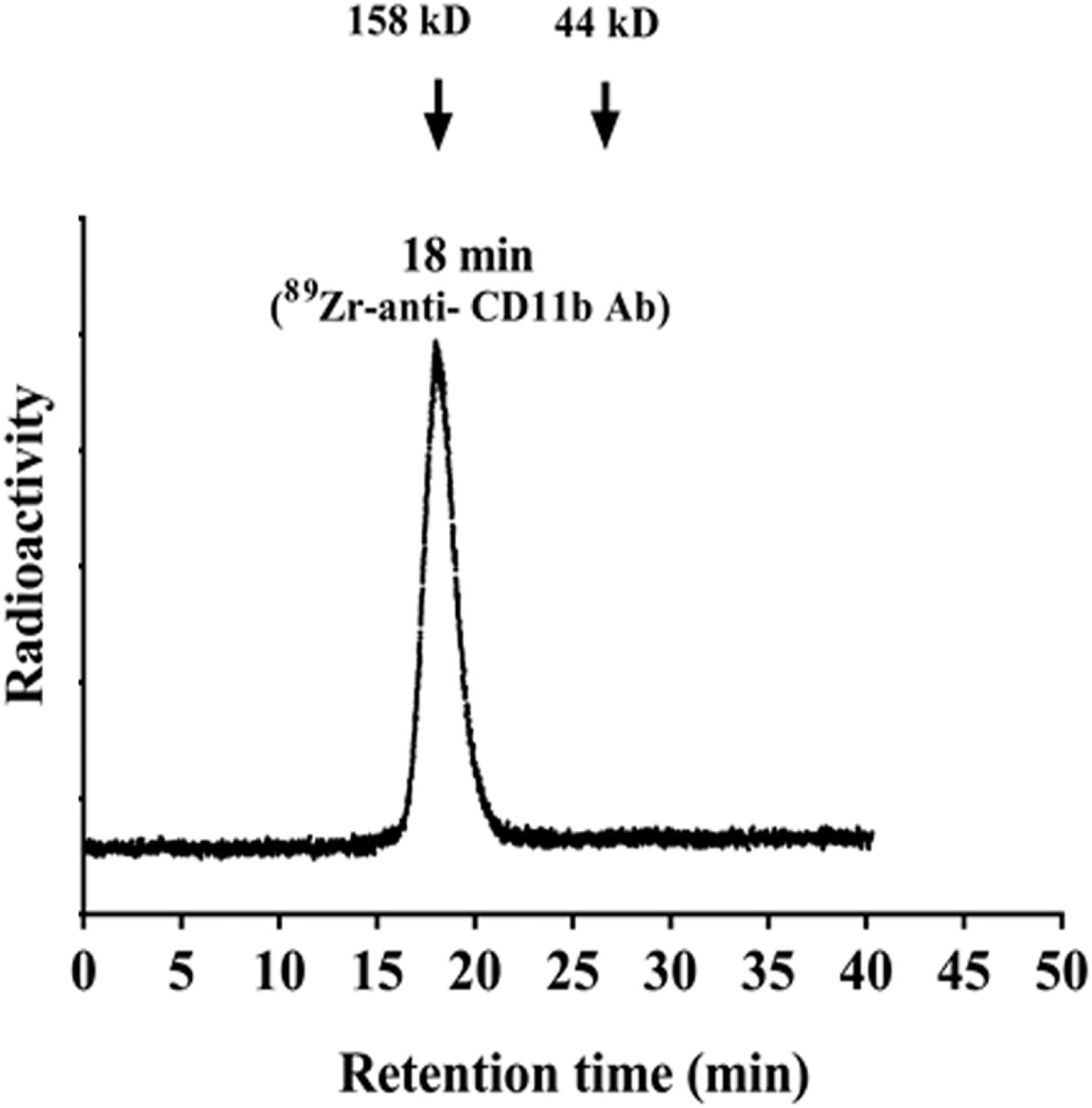
Radiochemical yield and purity of ^89^Zr-anti CD11b. Radio-SEC-HPLC chromatogram of the ^89^Zr-anti-CD11b (retention time 18 min), molecular weight marker showed 18.2 min retention time for 158 kD and 26.2 min retention time for 44 kD in the UV chromatogram.

### ImmunoPET imaging and biodistribution analysis

To corroborate the ability of ^89^Zr-anti-CD11b to detect CD11b^+^ cells in an *in vivo* GBM model, immunoPET was performed on mice bearing GL261 syngeneic gliomas 72 h post-injection of ^89^Zr-anti-CD11b Ab **(Fig. 2a-c)**. The experimental and blocking groups consisted of 6 mice each, with the latter group receiving radiotracer with a 10-fold lower specific activity as a block. All 6 experimental mice had greater uptake in the tumor ipsilateral right brain (SUVmean ± SD = 2.57 ± 0.25) than the contralateral left brain (SUVmean ± SD = 0.61 ± 0.11; p< 0.001) **(Fig. 2a-2d)**. High tracer uptake occurred the in spleen and lymph nodes, while muscle uptake was low **(Fig. 2c, 2e)**. Blocking with 10-fold lower specific activity of ^89^Zr-anti-CD11b Ab (**Fig. 2b, 2e**) resulted in significantly lower tumor ipsilateral right brain uptake (SUVmean ± SD = 0.12 ± 0.06) than for experimental mice (p<0.001) **(Fig. 2e)**. Average SUVmean was also higher in experimental mice for contralateral left brain, heart, and the right/left brain ratio (adjusted p<0.001) **(Fig. 2d)** and bone (adjusted p=0.04) but not for liver or muscle (adjusted p>0.42)

**Figure 2.**
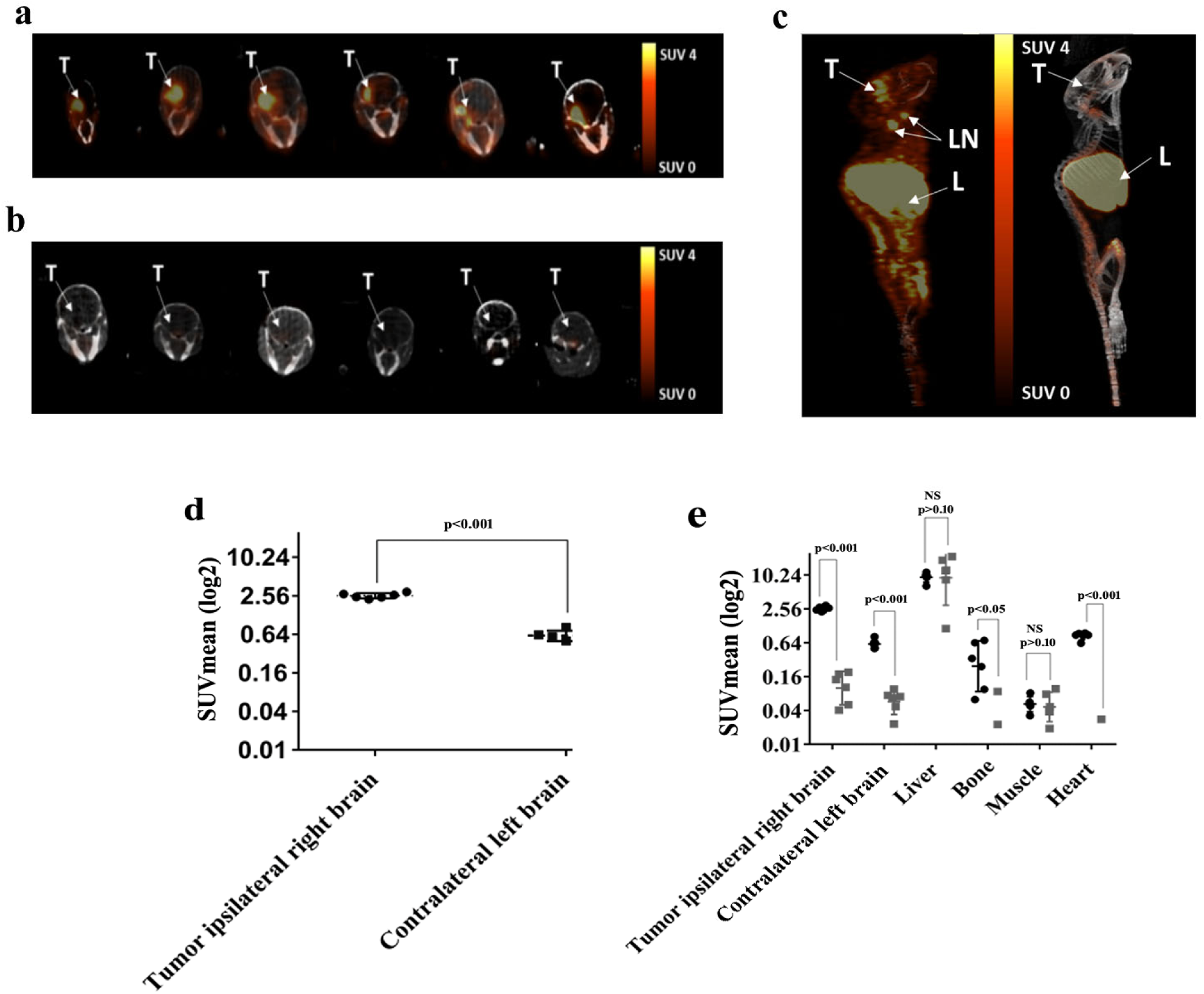
PET imaging of GL261 tumor-bearing mice with ^89^Zr-anti CD11b. ImmunoPET images and ROI analysis of GL261tumor-bearing mice. a) PET-CT axial images of the brain 72 h post injection of ^89^Zr-anti-CD11b Ab (n=6), tumor uptake is highly visualized in the ipsilateral right brain, SUVmean ± SD (2.57 ± 0.25). b) PET-CT axial images of the ipsilateral right brain 72 h post injection with 10-fold lower specific activity blocking dose (n=6), SUVmean ± SD (0.12 ± 0.06). c) PET-CT maximum intensity projection images of experimental (left) and control mice (right). T: tumor, LN: lymph node, L: liver. d) SUVmean of tumor ipsilateral right brain (•) and contralateral left brain for experimental mice (▪), p<0.001 (paired t-test). e) log2-transformed PET SUVmean values of major organs (p-values comparing experimental (•) and blocked (▪) are Benjamini-Hochberg adjusted). SUVmean values <0.01 (4 bone, 5 heart) not shown. Each plot shows mean ± SD, as well as individual data points.

Animals were euthanized after PET imaging, and major organs were collected for biodistribution studies to complement the *in vivo* results. Higher ^89^Zr-anti-CD11b Ab uptake was observed in blood, lung, muscle, heart, intestine, and thymus of the experimental group compared with the blocked group, but the differences were not statistically significant **(Fig. 3a)**. Uptake in the right brain, which contained the tumor, was significantly reduced by blocking with 10-fold decrease in specific activity; p = 0.028 after adjusted for multiple comparisons. Biodistribution uptake values showed the same pattern as the PET SUVmean uptake with CD11b in the tumor ipsilateral right brain (8.62 ± 3.29 %ID/g) was greater relative to the paired contralateral left brain (1.22 ± 0.09 %ID/g) (**Fig. 3b**, p<0.001). Tumor:blood and tumor:muscle ratios were relatively high (32.84 ± 66.15 and 26.72 ± 36.2, respectively). The high standard deviation values were due to very low muscle and blood % ID/g uptake in some mice.

**Figure 3.**
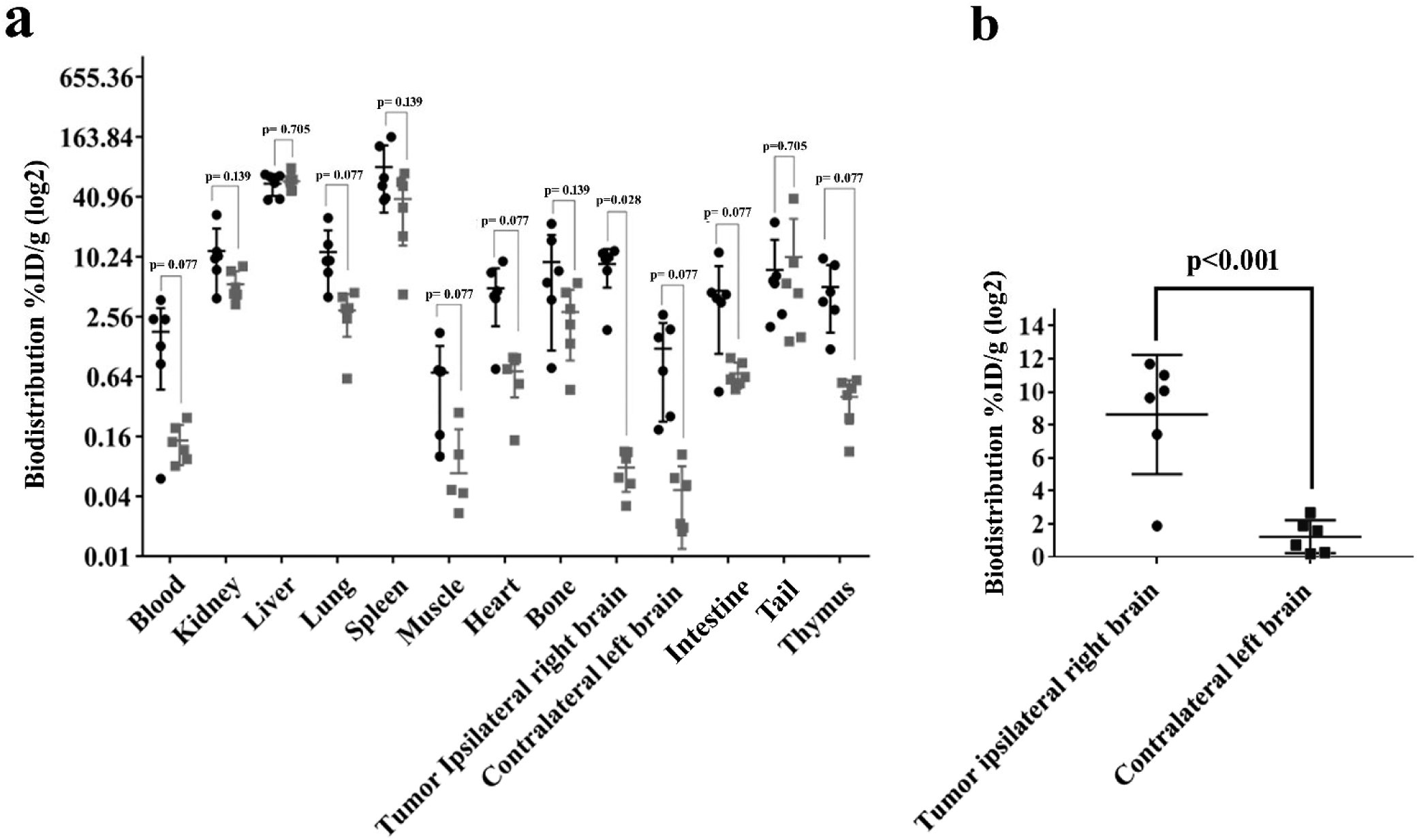
*Ex vivo* biodistribution of ^89^Zr-anti-CD11b in GL261 syngeneic glioma-bearing mice. Biodistribution analysis was performed 72 h post radiotracer injection. a) log2-transformed %ID/g values for major organs (p-values comparing experimental (•) and blocked (▪) are Benjamini-Hochberg adjusted). b) Biodistribution of paired tumor ipsilateral right brain (•) and contralateral left brain in experimental mice (▃), p<0.001. Each plot shows mean ± SD, as well as individual %ID/g data points.

### Immunostaining validation and receptor density of CD11b

To validate the presence of CD11b^+^ immune cells within the tumor and determine receptor density, GL261 tumors were dissected, dissociated, and analyzed by flow cytometry. Total immune cells were gated on using the forward scatter-area (FSC-A) and side scatter-area (SSC-A) parameters. TAMCs were identified by gating on CD45^+^CD11b^+^ cells (**Fig. 4a**, quadrant 2 [Q2]). The PE geometric mean of anti-CD11b on immune cells in GL261 tumor was determined to be 22,994 **(Fig. 4b)**, indicating an average of 54,076 CD11b receptors/cell.

**Figure 4.**
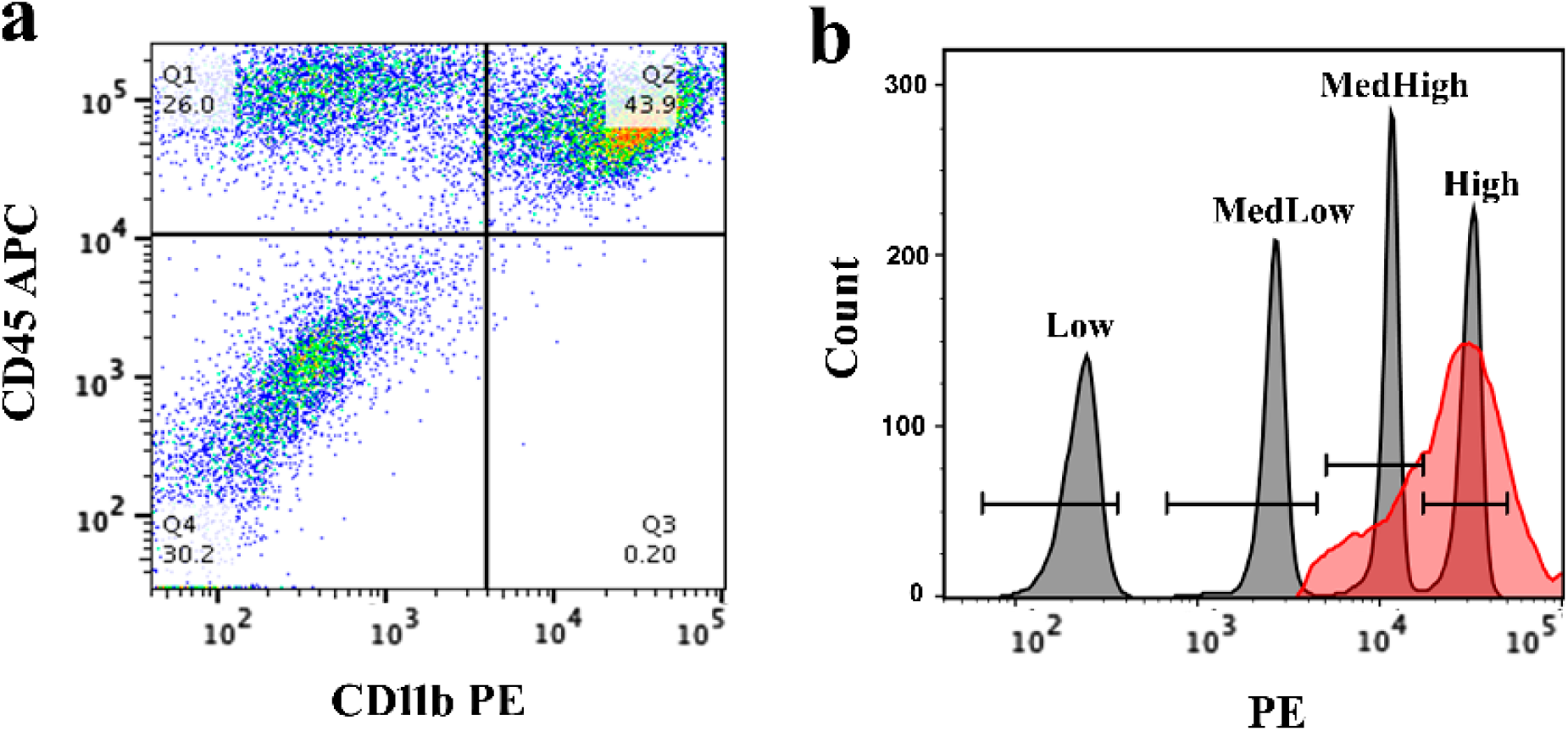
Quantification of CD11b on tumor-infiltrating immune cells. a) CD11b PE/CD45 APC dot plot, with Q2 containing CD45^+^CD11b^+^ TAMCs. b) histogram plot of QuantiBRITE PE bead singlets (gray peaks) and CD45^+^CD11b^+^ TAMCs (red peak). Bead population geometric means were determined to be 222 (Low), 2,528 (MedLow), 11,171 (MedHigh), and 30,585 (High). The CD45^+^CD11b^+^ TAMC geometric mean was found to be 22,994.

H&E staining was performed on the tumor sections from the right hemisphere and normal tissue from the contralateral left brain **(Fig. 5a-b)**. To determine the extent of intratumoral CD11b cell infiltration in the tumor relative to normal tissue, immunohistochemistry was performed. Contralateral brain and tumor sections were stained with anti-CD11b Ab (Clone EPR1344). Consistent with our PET and biodistribution analysis, tumors had an elevated number of CD11b^+^cells **(Fig. 5c, 5e)** compared with contralateral “normal” brain tissue **(Fig. 5d, 5f)**. ImageJ software (ImmunoRatio plugin) was used to analyze the percent of CD11b^+^ cells per nuclear area in IHC sections, which showed 62.3% ± 15.9% CD11b^+^ cells per nuclear area in the tumor tissue compared to 17.7% ± 10.9% in normal brain **(Fig. 5g-5k)**.

**Figure 5.**
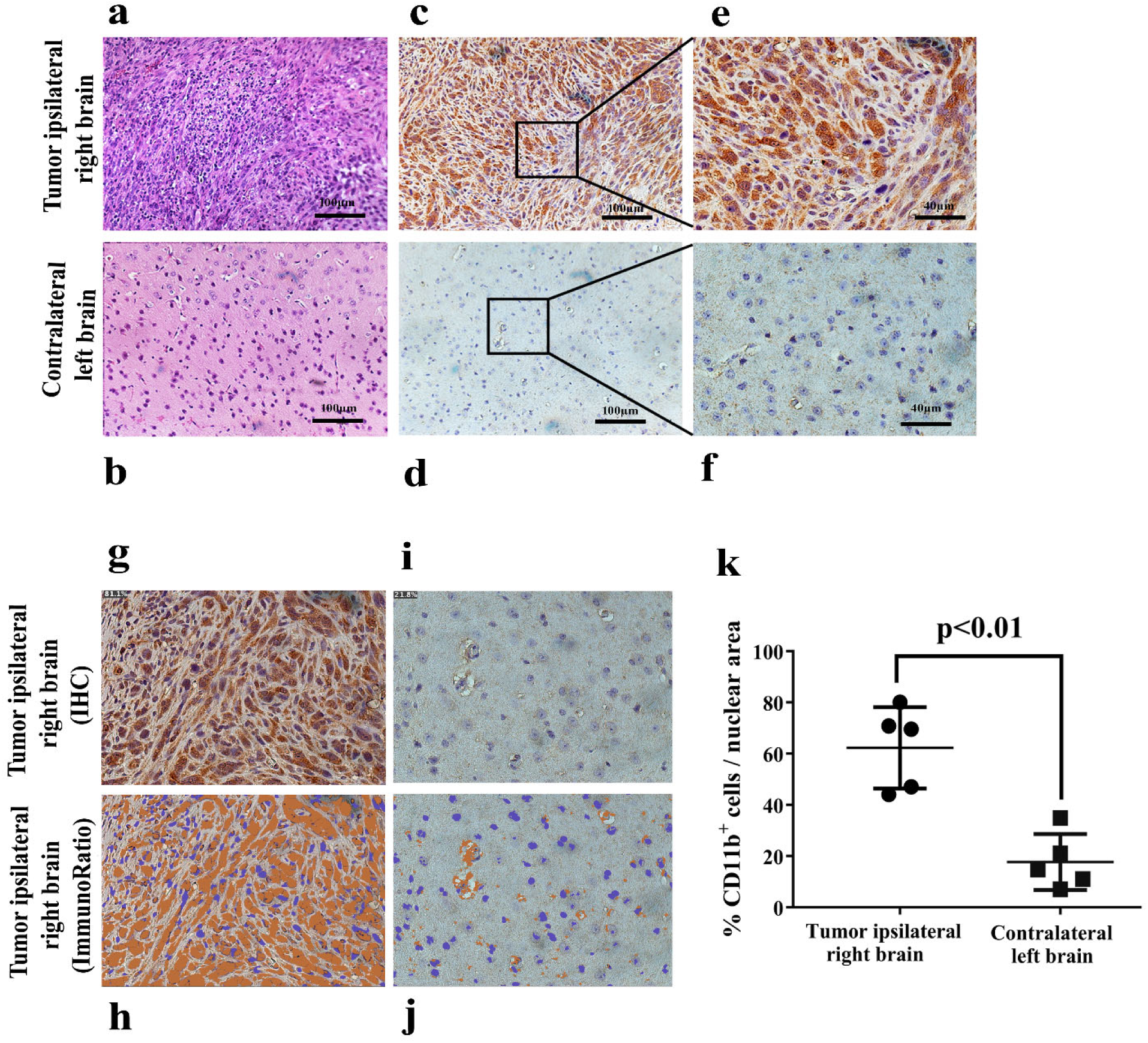
IHC validation and quantification of CD11b^+^ TAMCs in GL261 tumor. H&E and immunohistochemistry of the tumor from the ipsilateral right brain and normal tissue from the contralateral left brain. a) H&E of the tumor. b) H&E of normal tissue. c) Immunohistochemistry of the tumor at 20X magnification. d) Immunohistochemistry of normal tissue at 20X magnification. e) Immunohistochemistry of the tumor at 40X magnification. f) Immunohistochemistry of normal tissue at 40X magnification. g) Original IHC image of the tumor. h) Immunoratio of the tumor. i) Original IHC image of normal tissue. j) Immunoratio of normal tissue. k) Percent of CD11b^+^ cells per nuclear area of IHC images in the tumor ipsilateral right brain (•) and contralateral left brain in experimental mice (▃), p<0.01.

## Discussion

GBM is highly aggressive, immunosuppressive, and challenging to treat with current standard of care including surgery, chemotherapy, and radiation therapy [22]. In recent years, immunotherapy has emerged as a promising approach to enhance the efficacy of GBM therapies, either through boosting the immune system to attack the tumor or by eliminating immunosuppressive cells. The most abundant population of immune cells in GBM is the CD11b-positive TAMCs. TAMCs, primarily TAMs and MDSCs, may account for up to 40% of a GBM’s cellular mass. TAMCs are a promising clinical target due to their pro-tumoral and immunosuppressive features.

The relevance of TAMCs in promoting tumor growth in GBM was demonstrated by Rajappa et al., who investigated the role of CD11b^+^ cells in low- and high-grade murine gliomas. Administration of AZD1480, a JAK 1/2 inhibitor resulted in a significant decrease in tumor volume and prolonged survival *in vivo* by impairing CD11b^+^ cells recruitment to the tumor [23]. Additionally, numerous other preclinical studies have highlighted the tumor-promoting and immunosuppressive roles of these cells in GBM [2, 24-25]. However, unlike promising preclinical studies, efforts to deplete TAMCs in GBM patients have been disappointing. In a recent trial targeting TAMCs by inhibiting CSF-1R signaling with PLX3397, neither improvement in 6-month progression-free survival nor overall survival was observed compared with historical controls [26]. Although the median overall survival was 9.4 months, responses were varied, with overall survival times of up to ∼13 months. To understand these inconsistent outcomes, patient specimens were analyzed by IHC. Although no differences in intratumoral myeloid cells were found following PLX3397 therapy, PLX3397 did reduce chemotherapy-mediated recruitment of these cells into the tumor [26]. Due to the significant degree of heterogeneity in gliomas, the analysis of small tumor sections by IHC likely did not reflect the global number of these cells within patient tumors. Therefore, incorporating immunoPET into clinical trials to quantify TAMC populations could resolve the loss of temporal and spatial resolution associated with tissue sampling and enable the correlation of TAMC levels with therapeutic efficacy. In addition to PLX3397, other agents are being explored to inhibit TAMs and MDSCs [27]. For these reasons, we have pursued the development of an immunoPET tracer to quantify TAMs and MDSCs, or collectively, TAMCs.

TAMs and MDSCs share a common myeloid progenitor cell lineage, representing mature and undifferentiated developmental states, respectively. In both humans and mice, TAMs are typically identified as CD11b^+^CD14^+^ cells. While MDSC markers differ in mice and humans as CD11b^+^Gr1^+^ and CD11b^+^CD33^+^HLA-DR^-/low^, respectively, CD11b is common between the two. We have identified an anti-CD11b monoclonal antibody, clone M1/70, for preclinical evaluation as an immunoPET radiotracer in an orthotopic glioma model.

The syngeneic orthotopic GL261 mouse glioma model was used in this study. GL261 is the most extensively used syngeneic mouse glioma model for the preclinical study of glioma immunotherapy [28]. GL261 tumors exhibit profound anti-tumor immunity. Prolonged survival of mice following TAMC-targeted [29], checkpoint inhibitor [30-31], and vaccine immunotherapies [32-33] has supported the development of multiple clinical trial approaches for GBM patients [28].

To demonstrate the feasibility of immunoPET for quantification of tumor-infiltrating CD11b^+^ TAMCs, we administered ^89^Zr-DFO-anti-CD11b to mice bearing GL261 gliomas in the right hemisphere, while another group received the tracer at a blocking dose with 10-fold lower specific activity. CD11b^+^ cells were clearly visualized in the experimental group, and uptake was significantly reduced with excess unlabeled antibody, demonstrating CD11b-mediated accumulation of the radiotracer. Flow cytometry of dissociated GL261 tumor cells stained with anti-CD11b (clone M1/70) confirmed the presence of CD11b^+^ TAMCs and high levels of CD11b. These results demonstrate that TAMCs infiltrate gliomas in suitable numbers for visualization in preclinical PET. In addition to the dramatic uptake of ^89^Zr-anti-CD11b Ab at the tumor site compared to the contralateral left brain, the latter showed low levels of CD11b-mediated uptake, consistent with brain resident CD11b^+^ microglia being present in the intact uninflamed brain [34]. Marked uptake was also observed myeloid cell rich organs including spleen, bone, and thymus. High uptake in these organs was previously observed by Rashidian et al. with a radiolabeled anti-CD11b single domain antibody, where uptake corresponded to a significant number of myeloid cells within non-CNS tumors [35].

In conclusion, the data presented demonstrate that ^89^Zr-anti-CD11b Ab enables the surveillance of immunosuppressive TAMCs in the tumor microenvironment of a mouse glioma model, thereby providing a means of assessing the efficacy of therapeutic approaches. We demonstrated that CD11b is a highly promising imaging target for immunoPET based on sufficient CD11b receptor density, high frequency of CD11b^+^ TAMCs in gliomas, and the feasibility of an antibody-based PET tracer to detect these cells. Further studies are warranted to determine the effectiveness of CD11b immunoPET for determining differences in the infiltrative status of TAMCs before, during, and after therapy.

## Acknowledgments

We thank Kathryn Day and Joseph Latoche for assisting with preclinical PET/CT imaging and UPMC Hillman Cancer Center Histology Core for providing IHC assistance. This work was funded by National Institute of Health grant R21 EB026675 (WBE, GK), UPMC Hillman Cancer Center Animal Facility, In Vivo Imaging Facility, Biostatistics Facility (NCI P30 CA047904), and the St. Baldrick’s Foundation: for Pediatric Cancer Research, a St. Baldrick’s Hero Fund. No other potential conflict of interest relevant to this article is reported.

## Compliance with Ethical Standards

All applicable institutional and/or national guidelines for the care and use of animals were followed.

